# Relative Blindsight Induced by Incongruency with Working Memory

**DOI:** 10.1101/2025.11.14.688399

**Authors:** Wonyi Che, Hakwan Lau, Tomoya Nakamura

**Affiliations:** Center for Neuroscience Imaging Research, Institute for Basic Science, Suwon, Republic of Korea; Department of Brain Science and Engineering, Sungkyunkwan University, Suwon, Republic of Korea; Department of Intelligent Precision Healthcare Convergence, Sungkyunkwan University, Suwon, Republic of Korea; Department of Biomedical Engineering, Sungkyunkwan University, Suwon, Republic of Korea; Center for Brain Science, RIKEN, Wako, Japan; Japan Society for the Promotion of Science, Tokyo, Japan

## Abstract

The present study demonstrates a blindsight-like dissociation between subjective perception and general perceptual capacity in nonclinical observers. Participants performed either a two-interval forced-choice (2-IFC) or a Yes/No face detection task during working memory delay. To match general perceptual processing capacity across conditions, we titrated stimulus luminance contrast in the 2-IFC task and applied the same contrast in the Yes/No task. In Experiment 1, categorically incongruent (i.e., scene) information held in working memory impaired the sensitivity of the Yes/No detection of stimulus presence (face) vs. absence (blank background). In contrast, this effect was absent in Experiment 2, in which the Yes/No task required a discrimination of a face (target) vs. a scrambled face (non-target). These findings indicate that stimulus incongruency in working memory selectively influences subjective perception, dissociating it from general perceptual capacity.

## Introduction

Working memory and perception have been traditionally considered distinct cognitive processes. However, recent neuroscientific studies suggest they engage in overlapping neural representation. For example, using fMRI multivoxel pattern analyses, Harrison and Tong (2009) demonstrated that orientation representations maintained in working memory could be decoded from the early visual cortex using a classifier trained on perceptual responses (see also Albers et al., 2013; Rademaker et al., 2019). Similar cross-decoding has since been demonstrated in association cortices, particularly the prefrontal cortex, suggesting that perceptual and mnemonic representations share a common neural code even beyond early sensory areas (e.g., Mendoza-Halliday & Martinez-Trujillo, 2017; Nakamura et al., 2026).

Because of the overlap of their neural representation, actively maintained information in working memory may directly influence perceptual processing. Gayet et al. (2017), for example, have shown that maintaining the working memory contents enhances neural responses in the visual cortex when subsequent sensory input matches the mnemonic representation (a congruent stimulus).

In a similar vein, we may expect that working memory would interact with the perception of stimuli at the behavioral level. Indeed, previous studies (Kang et al., 2011; Teng & Kravitz, 2019) have shown that visual working memory can bias subsequent perception. For example, Kang et al. (2011) demonstrated that participants who maintained a certain dot motion direction in working memory perceived subsequent motion stimuli as shifted away from the remembered direction, which we can describe as a repulsion effect. Similarly, Teng and Kravitz (2019) suggested that similarity between the content of the working memory and the perception (e.g., color and orientation) systematically increased the degree of perceptual bias.

However, it is unknown whether working memory mainly influences the general capacity for distinguishing between different stimuli or whether it also influences the subjective experience of seeing. According to some theoretical perspectives, subjective experience may require some putative central mechanisms to distinguish internally generated representations (i.e., working memory and imagery) from external sensory inputs (Lau, 2019; Dijkstra & Fleming, 2023). If this is the case, working memory may influence subjective aspects of perception, even independently of the general capacity of perceptual processing.

Here, we tested this possibility by conducting psychophysical experiments, in which perceptual sensitivity was compared between a two-interval forced-choice (2-IFC) and a Yes/No (Y/N) task performed during the working memory delay period. In addition, to see the systematic effect of similarity (Teng & Kravitz, 2019), we also manipulated the congruence of working memory and perceptual content.

The 2-IFC performance is a standard measure of objective processing capacity. Since it depends on one’s ability to discriminate between a pair of external stimuli, the judgment is less likely to be influenced by subjective decision criteria (Macmillan & Creelman, 1991; but see Yeshurun et al., 2008). In contrast, the Y/N task can be considered a relatively subjective task, as it requires a certain internal template or reference to compare against the single stimulus (Michel & Lau, 2021). Therefore, capturing variation in Y/N task performance while holding 2-IFC performance constant provides one way to isolate changes in subjective perception (Samaha, 2015).

Conceptually, such a distinction resembles blindsight, a neurological condition in which patients damaged in their primary visual cortex deny seeing stimuli presented in their damaged visual field while showing above-chance performance in forced-choice discrimination tasks (Weiskrantz et al., 1974). Importantly, Azzopardi and Cowey (1997) found that Y/N sensitivity relative to 2-IFC was impaired in blindsight, consistent with the idea that Y/N sensitivity may selectively track subjective (i.e., conscious) perception, at least in ways more than 2-IFC sensitivity does.

Essentially, the present study tested whether manipulating working memory congruence could produce a blindsight-like dissociation between subjective and objective measures of perception in healthy observers. We investigated how working memory congruence influences Y/N performance under two different perceptual task designs. In Experiment 1, we used a pure detection paradigm in which the task required the judgment of the stimulus presence/absence, finding that Y/N sensitivity dropped only when the participants had to hold incongruent content in working memory. However, such selective modulation in Y/N performance disappeared in Experiment 2, in which the task required discriminating a face from a scrambled face.

## Methods

The experiments are presented in an order that best supports the scientific narrative rather than the chronological order in which they were conducted. The study began with a Preliminary experiment (reported in the Supporting Information), in which data were initially collected from eight participants. Based on these preliminary data, we performed a power analysis and preregistered the design and analysis plan for Experiment 2, which was then conducted with a new sample of 16 participants. To match the statistical power of the preliminary data, an additional eight new participants were then recruited for the Preliminary experiment, bringing its final sample size to 16. Finally, Experiment 1 (presented first in the main text) was conducted with another independent sample of 16 participants. This reordering is intended to improve the clarity of presentation and does not affect the interpretation and conclusions of the results.

### Experiment 1

#### Participants

A total of 16 university students (8 females, 8 males, mean age = 22.6 years, *SD* = 4.3) participated in Experiment 1. All were naïve to the purpose of the study and self-reported normal or corrected-to-normal visual acuity, no color vision deficiency, and no prior psychiatric diagnosis. All participants provided written informed consent under procedures approved by the Institutional Review Board of Sungkyunkwan University.

#### Apparatus

The experiment was programmed and run using PsychoPy version 2024.2.4 (Peirce et al., 2019). Visual stimuli were presented on an HP p1230 gamma-calibrated CRT monitor with a resolution of 1024 × 768 pixels at a refresh rate of 60 Hz (width: 39.37 cm, corresponding to a 900-pixel-wide display for the stimulus, 8-bit color depth, standard dynamic range). A CRT monitor was chosen because of its high temporal precision and minimal latency, characteristics that make it well-suited for psychophysical experiments. In a darkened room, participants viewed the stimuli on the monitor at a fixed distance of 54.1 cm, with their head stabilized on the head-and-chin rest.

#### Stimuli

As visual stimuli, we used faces, scenes, masks, and scrambled faces, all presented to participants in grayscale, on a uniform gray background (21 cd/m²). The experiment included three working memory/perception congruence conditions: stimulus-congruent (SC), category-congruent (CC), and category-incongruent (CI). In the SC condition, face perceptual stimuli were physically identical to a face used in the working memory contents; in the CC condition, face perceptual stimuli were drawn from different exemplars within the same face category; and in the CI condition, we used a different category of stimuli (scenes) from the face perceptual tasks. The face images were selected from the Chicago Face Database: A free stimulus set of faces and norming data (Ma et al., 2015; http://www.chicagofaces.org), and the scene images were from the *Scene Size x Clutter Database* (Park et al., 2015). We used a pixelwise random noise image (generated and saved in advance) as a mask. We generated scrambled face stimuli by applying the Fourier image processing phase, which scrambles the original face images into quadrants and rearranges them. Original face images were cropped to remove background regions. The two-dimensional Fast Fourier Transform (FFT) of each image was then computed, and the phase spectrum was divided into four quadrants. Within each of the first and second quadrants, the phase values were randomly permuted along both the horizontal and vertical axes. The third and fourth quadrants were obtained by reflecting the permuted second and first quadrants across the horizontal axis, respectively. The permuted phase matrices were recombined with the original amplitude spectrum and inverse Fourier-transformed (IFFT) to reconstruct the scrambled images. This procedure preserves the global amplitude (i.e., spatial frequency and luminance distribution) of the original faces while disrupting configural information. The scrambled faces, therefore, served as controls, ensuring that low-level visual properties were matched while facial identity was removed.

In the 2-IFC tasks, one of two intervals contained a face stimulus, and the other contained a blank. In the Y/N task, either a face or a blank was present. Both perceptual tasks occurred during the delay of the working memory task, and the trials were equally distributed across three different congruencies (Stimulus Congruent: SC, Category Congruent: CC, Category Incongruent: CI; see Fig. 1 for further details). Congruence was defined by whether the perceptual stimuli matched the content of the working memory task. In the congruent conditions, participants memorized faces, whereas in the category incongruent condition, they memorized scenes, while the perceptual tasks involved faces. We included both stimulus- and category-congruent conditions to establish a gradient of similarity between the working memory content and the perceptual input, allowing us to distinguish image-level (SC) from broader category-level (CC) influences on perceptual decisions.

The sample and probe for the working memory task were presented at four isoeccentric peripheral locations (11.7°). We asked participants to maintain their gaze on a fixation point displayed at the center of the screen during the 500-ms delay before perceptual stimulus onset. The face stimuli for the 2-IFC and Y/N tasks were presented at fixation. In each experimental block, the same sample stimuli were shown only once, except in the case of the SC condition. Sample stimuli were randomised during each trial and for each set of runs.

To match performance (∼79%) across congruence conditions, the root-mean-square (RMS) contrast of the face stimuli in the 2-IFC task was adaptively controlled using a 1-up-3-down staircase procedure (Levitt, 1971), separately for each of SC, CC, and CI conditions. The initial contrast value was 2.3%, and the values were bounded between 0.005% and 14.1% on a logarithmic scale to avoid reaching zero contrast. Each staircase consisted of 33 trials. The contrast value from the last (33rd) trial was taken as the final converged threshold, which was then used as the fixed contrast in the subsequent Y/N task under the same congruence condition.

#### Procedure

We used a 2 (perceptual tasks: 2-IFC, Y/N) × 3 (congruence: SC, CC, CI) within-subject factorial design. The experiment was conducted over two days, with three blocks (approximately 1.5 h) completed per day. Each block consisted of two sub-blocks corresponding to the two perceptual tasks: the first sub-block was for the 2-IFC task, followed by the second sub-block for the Y/N task. Within each sub-block, 33 trials from each of three conditions (SC, CC, CI) were performed in random order. Participants were allowed to take an optional 5-minute break between blocks. In total, participants completed six such blocks, yielding 1,188 trials (6 blocks × 33 trials × 2 perceptual tasks × 3 congruence conditions).

On the first day, participants read the task instructions and completed 15 practice trials in the CC condition (3 trials for each: working memory task only, 2-IFC task only, Y/N task only, combined working memory-2-IFC, and combined working memory-Y/N) with trial-to-trial feedback.

In the main experiment, participants were asked to perform either the 2-IFC or the Y/N task during the working memory delay period (Fig. 1). Experiment 1 used a pure detection design in which either a face stimulus or no stimulus (rather than a scrambled face) was presented in both the 2-IFC and Y/N tasks. To indicate the potential stimulus interval, a brief auditory cue was delivered at the expected stimulus onset, thereby matching temporal uncertainty across perceptual tasks. At the beginning of each trial, as working memory sample stimuli, four different images were presented for 2000-ms, followed by a 500-ms delay.

In Experiment 1, the 2-IFC sub-block, two stimuli (a face and a blank) were sequentially shown in a random order, for 200-ms each, with a 500-ms inter-stimulus interval. Following the 300-ms mask, participants had 2000-ms to indicate whether the face appeared in the first or second interval, using left or right arrow keys.

In the Y/N sub-block, either a face or just a blank was presented in a single interval, and participants judged whether a face appeared or not. Each face stimulus was followed by a 300-ms mask.

Regardless of the perceptual task, right after the response on the perceptual task, the working memory probe was presented: one of the four images was replaced from the initial sample sets, and the participants were asked to report which was the new image using the 1, 2, 3, or 4 keys within the time limit of 3000-ms. This probe display was followed by a 1500-ms inter-trial interval. A beep sound was played when they did not press any keys within the time limit. Nevertheless, the response was recorded up until the appearance of a working memory sample for the next trial, even after the beep sound. In the main experiment, no feedback was provided for either the working memory or the perception task.

### Experiment 2

#### Participants

Another 16 naïve university students (11 females, 5 males, mean age = 23.9 years, *SD* = 2.9) who were naïve to the purpose of the study participated in Experiment 2. The eligibility criteria and ethical procedures were identical to those in Experiment 1. The sample size, experimental procedures, and analysis plan were pre-registered (https://www.biorxiv.org/content/10.1101/2025.11.14.688399v3).

In the 2-IFC task, one of two intervals contained a face stimulus, and the other contained a scrambled image matched in contrast, luminance, and size. In the Y/N task, either a face or a scrambled face was presented.

#### Procedure

Experiment 2 was identical to Experiment 1 except that scrambled face stimuli were presented in both the 2-IFC and Y/N tasks. Experiment 2 better matches the stimulus, thereby enabling a fairer comparison of detection sensitivity across tasks. The apparatus and the stimuli were identical to those used in Experiment 1.

### Data Analysis

#### Sensitivity in perception tasks

For the perception task, we calculated detection sensitivity, i.e., participants’ ability to discriminate between faces (signals) and blank screens or scrambled faces (noise), depending on the experiment, according to signal detection theory (SDT; Macmillan & Creelman, 1991). All the following analyses were based on the unequal-variance SDT model because variances for the signal and noise distributions are usually different, especially for the Y/N task.

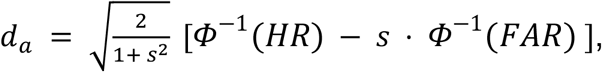

where *Φ*^−1^ denotes the z-transformation of hit rate (HR) and false alarm rate (FAR). For the Y/N task, HR was the proportion of “Yes” responses to the signal, and FAR was the proportion of “Yes” responses to the noise:

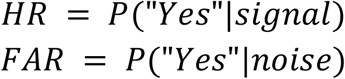

To calculate the HR and FAR for the 2-IFC task, we treated the signal in the first interval as the “signal”:

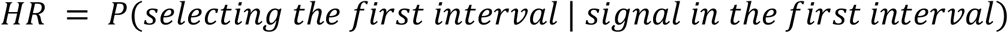

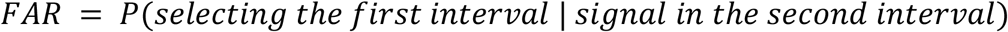

To avoid obtaining infinite z-values, 0 and 100% were adjusted by ± 0.5 prior to the z-transformation (Hautus, 1995). *s* represents the slope of the zROC, corresponding to the ratio of the standard deviations of the noise and signal distribution 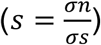. To get the estimate of *s*, we derived a type-1 zROC (the z-transformed receiver operating characteristic function) using reaction time (RT) data (Miyoshi et al., 2026). Capitalizing on the empirical observation that quicker RT is associated with higher confidence, we divided trials into three RT bins (fast, middle, slow) for each response type. By shifting the decision criterion across these bins, we obtained five pairs of HR and FAR. A linear function was then fitted to the resulting zROC, and *s* was estimated as the inverse of the slope of the best-fitting line.

#### Statistical test

To test our main hypothesis, we performed a one-way repeated-measures ANOVA on the sensitivity ratio across congruence conditions in all experiments. When the main effect was significant, we also performed Bonferroni-corrected two-tailed paired t-tests to examine pairwise differences (SC–CC, CC–CI, and SC–CI).

We performed follow-up repeated-measures ANOVAs for each task separately to characterize significant task-by-congruence interactions. HR, FAR, and staircase contrast thresholds were analyzed using one-way repeated-measures ANOVAs across congruence conditions, followed by Bonferroni-corrected paired t-tests when necessary. The assumption of sphericity was assessed using Mauchly’s test, and Greenhouse-Geisser corrections were applied when the assumption of sphericity was violated.

## Results

### Working memory modulation of Y/N pure detection under matched stimulus alternatives

We conducted Experiment 1 using a pure detection version of the Y/N task, in which participants had to judge whether the face stimulus was present (Yes) or absent (No), to examine whether working memory could selectively impair Y/N sensitivity when the task required subjective detection.

In Experiment 1, 16 participants performed a working memory task, in which they memorized four sample stimuli and, after a delay, selected which of four probe stimuli had changed relative to the sample (Fig. 1). During this working memory delay, they also performed a perceptual task: either a 2-IFC task or a Y/N task. In the 2-IFC task, the participants were asked to discriminate between a face and a blank presented in separate temporal intervals, whereas in the Y/N task, they were asked to judge whether a single interval contained a face or was blank (i.e., face pure detection against blank). The working memory samples were either four scenes (category-incongruent), four faces of different identities from the perceptual face stimulus (category-congruent), or four faces including the same identity as the perceptual face stimulus (stimulus-congruent).

**Figure 1.**
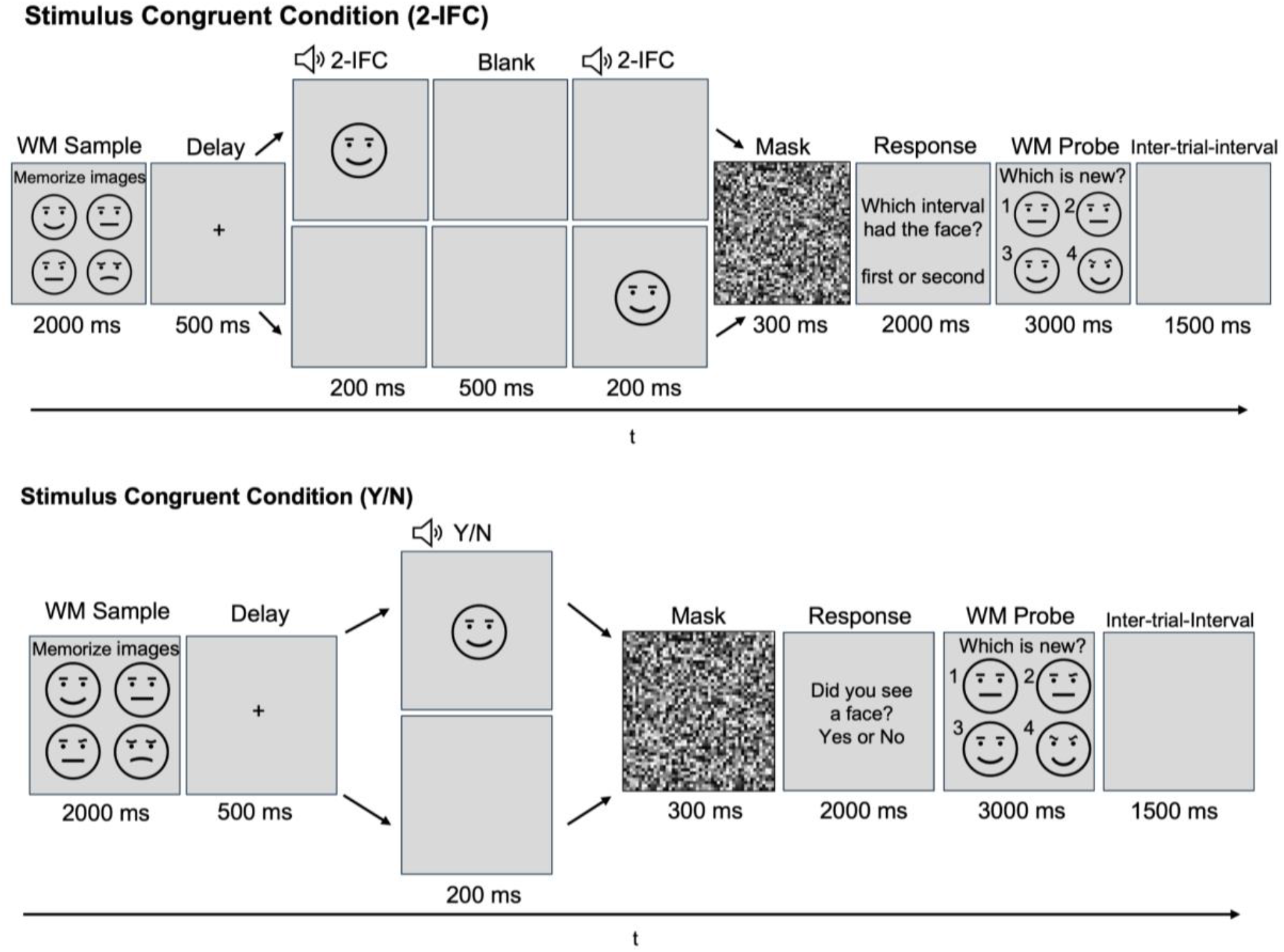
Experimental paradigm and task conditions. In the Stimulus Congruent (SC) condition, one of the working memory samples was physically identical to the face stimulus in the perceptual task (2-IFC and Y/N). In the Category Congruent (CC) condition, working memory stimuli belonged to the same category (faces) as the perceptual stimulus but were not physically identical. In the Category Incongruent (CI) condition, working memory stimuli belonged to a different category (scenes) from the perceptual face stimulus. Regardless of condition, a single trial consisted of a working memory sample display (encoding 4 images), a delay, a perceptual task (2-IFC or Y/N face detection), and finally, a working memory probe display in which participants identified the novel stimulus. For copyright reasons, face stimuli were substituted in this manuscript with copyright-free emoji exemplars, only for illustrative purposes. In Experiment 1, the perceptual task used a blank interval, accompanied by a brief auditory cue to indicate stimulus timing. In Experiment 2, we used a scrambled face, which was shown in all the perceptual tasks.

According to signal detection theory, sensitivity in a 2-IFC task is expected to exceed that in a Y/N task by a factor of √2 (Macmillan & Creelman, 1991; Azzopardi & Cowey, 1997):

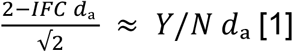

2-IFC sensitivity scaled by √2 was comparable to Y/N sensitivity (SC: scaled 2-IFC *d_a_ =* 1.101, Y/N *d_a_ =* 1.325; CC: scaled 2-IFC *d_a_ =* 1.142, Y/N *d_a_ =* 1.086; CI: scaled 2-IFC *d_a_ =* 1.058, Y/N *d_a_ =* 0.764). After scaling, there was no significant main effect of task (*F*(1,15) = 0.248, *p* = .626, 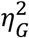 = .004), indicating the theoretical relationship between Y/N and 2-IFC sensitivity was preserved overall (Fig. 2a).

As our primary analysis, the detection sensitivity (*d_a_*) ratio showed a significant main effect of congruence, *F*(2,30) = 6.909, *p* = .003, 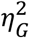 *=* .192. Pairwise comparisons showed that the ratio was significantly higher in the SC condition than in the CI condition, *t*(15) = 4.104, *p* < .001, Hedges’ *g* = 1.296. The SC–CC comparison also reached significance, *t*(15) = 2.195, *p* = .044, Hedges’ *g* = .529. However, the CC–CI comparison was not significant, *t*(15) = 1.532, *p* = .146, Hedges’ *g* = 0.536.

To see why Y/N sensitivity (*d_a_*) selectively drops in the CI condition, we compared hit rate (HR) and false alarm rate (FAR) separately across conditions (Fig. 2b). There was a significant main effect of congruence on both HR and FAR (*F*(2,30) = 30.094, *p* < .001, 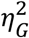 *=* .440; *F*(2,30) = 12.494, *p* < .001, 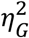 *=* .080). Pairwise comparison indicated that HR was significantly reduced in the CI condition relative to both SC and CC conditions (SC–CI: *t*(15) = 8.169, *p* < .001, Hedges’ *g* = 1.856, CC–CI: *t*(15) = 5.596, *p* < .001, Hedges’ *g* = 1.546). FAR also decreased significantly in the CI condition (SC–CI: *t*(15) = 3.812, *p* = .002, Hedges’ *g* = .484, CC–CI: *t*(15) = 3.922, *p* = .001, Hedges’ *g* = .656), but to a smaller extent than HR. Therefore, the decrease in Y/N sensitivity in the CI condition was primarily driven by a failure to notice the stimulus.

Contrast thresholds used in the Y/N task were significantly different across conditions (*F*(2,30) = 13.016, *p* <.001, 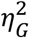 *=* .093) (Fig. 2c). Additionally, the CI condition required significantly higher contrast values than both the SC and CC conditions (SC–CI: *t*(15) = -4.655, *p* <.001, Hedges’ *g* = -0.666; CC–CI: *t*(15) = -3.615, *p = .002,* Hedges’ *g* = -0.576), whereas no significant difference was observed between the SC and CC conditions (SC–CC: *t*(15) = -0.347, *p = 0.733,* Hedges’ *g* = -0.044). Thus, the sensitivity decrease observed in the CI condition cannot be explained simply by the higher stimulus intensity used in that condition, as increased stimulus intensity would be expected to produce the opposite effect.

**Figure 2.**
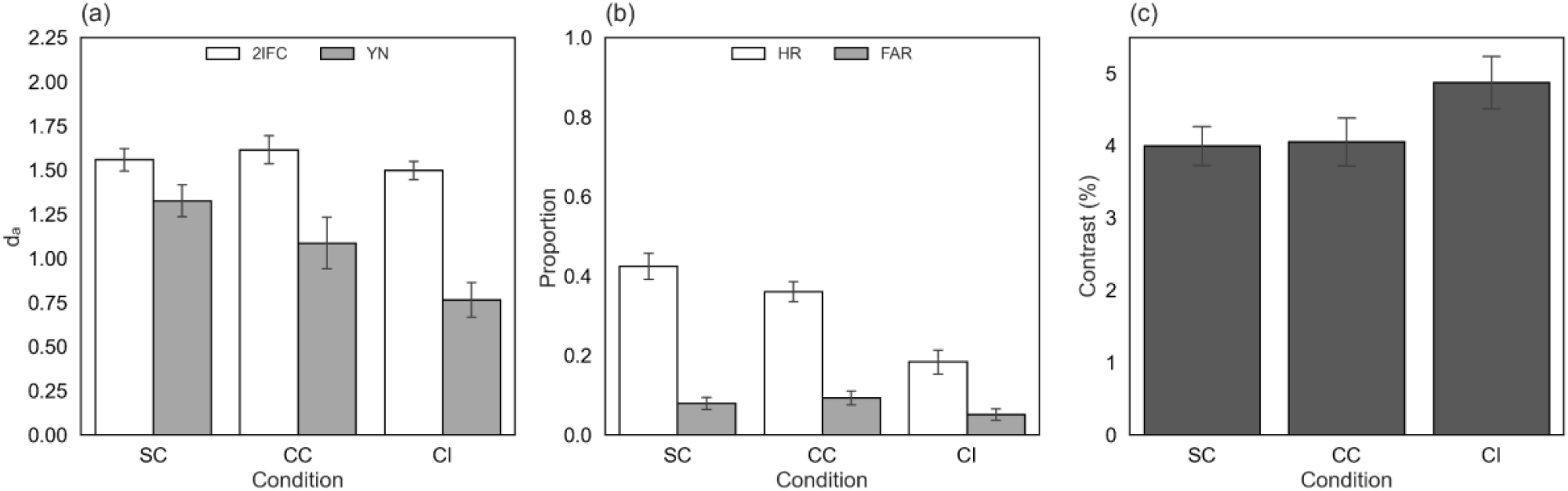
Results of Experiment 1. (a) Sensitivity (dₐ) in the 2-IFC and Y/N tasks. (b) Hit Rate (HR) and False Alarm Rate (FAR) in the Y/N task. (c) Contrast thresholds derived from staircases in 2-IFC tasks. Bar graphs indicate inter-individual averages, with error bars indicating standard errors of the mean.

### Working memory did not modulate the Y/N discrimination performance

In Experiment 2, 16 newly recruited participants performed either a 2-IFC task or a Y/N task during the working memory delay, as in Experiment 1. Unlike in Experiment 1, a blank interval was replaced with a scrambled face in the Y/N task, ensuring that the stimulus alternatives were fully matched with the 2-IFC task (i.e., face vs scrambled face judgments).

The theoretical relationship between Y/N and 2-IFC sensitivities was also preserved in Experiment 2 (Fig. 3a). 2-IFC sensitivity scaled by √2 was comparable to Y/N sensitivity (SC: scaled 2-IFC *d_a_ =* 1.078, Y/N *d_a_ =* 1.100; CC: scaled 2-IFC *d_a_ =* 1.122, Y/N *d_a_ =* 1.098; CI: scaled 2-IFC *d_a_ =* 1.028, Y/N *d_a_ =* 0.942), which was statistically supported by no significant effect of task (*F*(1,15) = 0.183, *p* = .675, 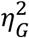 = .002).

Unlike in Experiment 1, the detection sensitivity ratio revealed that the main effect of congruence was not significant (*F*(2,30) = 0.172, *p* = .842, 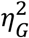 *= .007*). Therefore, we found no evidence that working memory could selectively modulate Y/N sensitivity when the task required discriminating faces from another stimulus (i.e., a scrambled face).

HR and FAR were also comparable across conditions in Experiment 2 (Fig. 3b). There was no main effect of congruence on either HR (*F*(2,30) = .484, *p* =.621, 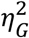 =.007) or FAR (*F*(2,30) = .071, *p* =.932, 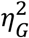 =.000).

The contrast threshold (i.e., stimulus intensity used in the Y/N task) was also comparable across conditions (*F*(2,30) = 0.378, *p* = .689, 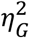 *=* .*008*).

**Figure 3.**
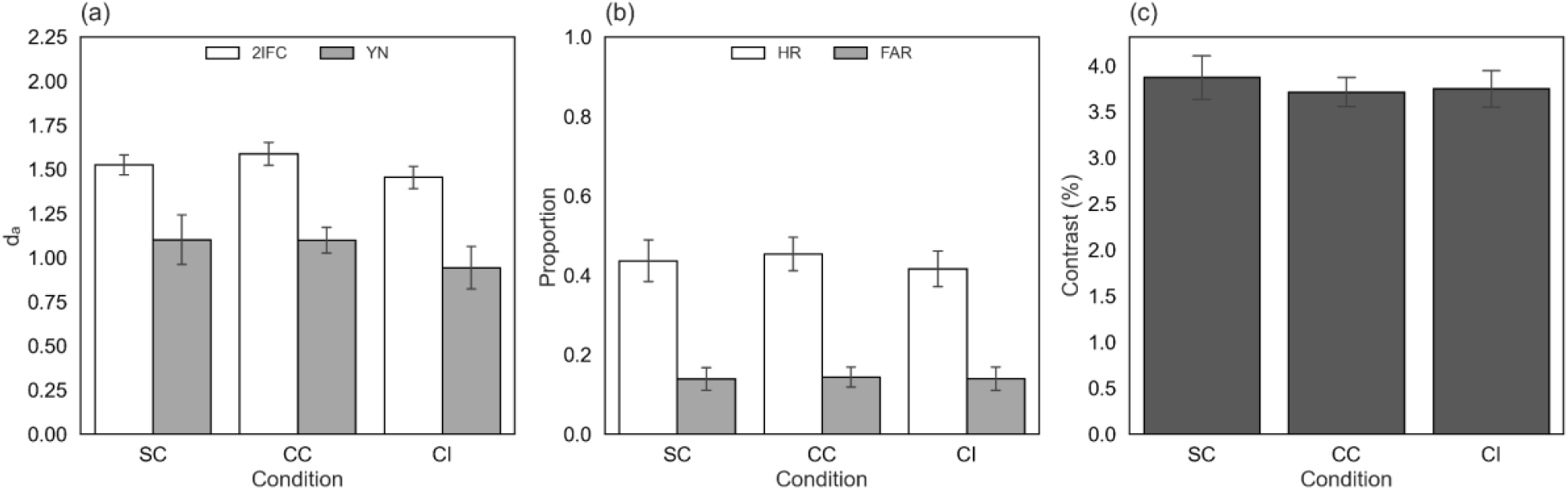
Results of Experiment 2. (a) Sensitivity (dₐ) in the 2-IFC and Y/N tasks. (b) Hit Rate (HR) and False Alarm Rate (FAR) in the Y/N task. (c) Contrast thresholds derived from staircases in 2-IFC tasks. Bar graphs indicate inter-individual averages, with error bars indicating standard errors of the mean.

## Discussion

In the present study, we found that working memory selectively impaired Y/N sensitivity under category-incongruent conditions despite matched general perceptual capacities as measured in a 2-IFC task. Importantly, this effect was observed only when the Y/N task required “pure detection” (Experiment 1), while it was absent when the Y/N task required discrimination between a face and a scrambled face (Experiment 2). Results from an earlier preliminary experiment also provided converging evidence for this selective effect on Y/N detection sensitivity (Fig S1,3).

In Experiment 2, the judgement between competing stimulus alternatives (face vs scrambled face) may also be thought of as a detection task, in that a face could be rather naturally construed as a target, and the scrambled face as a non-target.

However, the luminance contrasts for the two possible stimulus alternatives were identical. Whereas in the other experiments concerning what we call “pure detection”, the non-target was just blank, with zero stimulus energy. In this sense, the absence of the target was not a well-defined stimulus. The effect of blindsight-like dissociation here seems to be specific to this type of “pure detection” of stimulus vs its absence.

Consistent with signal detection theory, both experiments showed lower Y/N performance (accuracy and sensitivity) relative to 2-IFC performance. This is natural because the 2-IFC task provides twice as many opportunities as the Y/N task to view the stimuli that are informative for perceptual decisions (Macmillan & Creelman, 1991). Theoretically, the 2-IFC *d_a_* is expected to be √2 times the Y/N *d_a_*, as long as all factors are matched except for the number of stimulus presentations (Macmillan & Creelman, 1991; but see Yeshurun et al., 2008). In our data, the canonical √2 relationship was overall preserved except for the category incongruent condition in Experiment 1, where the critical experimental effect (i.e., selective modulation of subjective detection performance) was observed.

One possible explanation for the sensitivity reduction in the Y/N task is that the Y/N task depends more heavily on internal criterion-setting than 2-IFC tasks. In the 2-IFC task, participants can base their decision on the difference between two external stimulus alternatives. It is relatively intuitive to see what a zero difference would mean. In contrast, a Y/N task requires participants to evaluate a single external stimulus relative to internal noise, in such a way that optimal criterion placement may be less obvious (Ko & Lau, 2012; Michel & Lau, 2021). This may be a reason why the Y/N task, especially the “pure detection” variant, is particularly susceptible to cognitive manipulations that target internally maintained representations (e.g., incongruency with working memory content).

Another possibility is that the category incongruent condition may have incurred a cost of shifting attention away from faces. In this condition, participants first had to hold four different scene images while performing a face perception task. Before performing the perceptual task, the participants may have had to disengage from the maintained scene representation and redirect attention to external face stimuli.

Because working memory maintenance is a form of internal attention that competes with external perceptual selection (e.g., Kiyonaga & Egner, 2013), shifting attention between them can incur a cost to performance (e.g., Nobre & Gresch, 2025).

Moreover, shifting attention between different visual categories has been shown to incur additional switching costs (Walther & Fei-Fei, 2007). Although this evidence concerns switching between externally presented stimuli, it raises the possibility that switching between internally maintained and externally perceived representations from different categories may also impose additional attentional demands.

However, this account alone is unlikely to entirely explain our results. If attentional switching primarily imposed a general processing cost, it would be expected to affect perceptual performance irrespective of whether the task required “pure detection”, Y/N discrimination, or 2-IFC. However, the present study found a selective impairment only for “pure detection”.

The selective influence of working memory on subjective aspects of perception resembles the phenomenon of blindsight, in which patients with damage to the primary visual cortex can perform above chance on forced-choice visual tasks despite the absence of subjective reports of conscious experience. Similarly to our findings, Azzopardi and Cowey (1997) demonstrated that blindsight patients exhibit lower Y/N detection sensitivity than would be predicted from their forced-choice performance under standard signal detection theory assumptions (Equation 1).

Interestingly, a recent study in mice (Bhatla et al., 2026) provided some causal evidence of this dissociation. The inactivation of the dorsal hippocampus produced a robust blindsight-like behavior (Bhatla et al., 2026). This suggests that modulation of late-stage processing, especially in memory-related circuits, can selectively disrupt putatively conscious visual detection while relatively sparing some residual visual functions. Likewise, disrupting prefrontal function using transcranial magnetic stimulation has been reported to selectively impair metacognitive sensitivity while leaving the perceptual sensitivity intact (Rounis et al., 2010). Together, these findings suggest that blindsight-like dissociation may not happen only due to the lesions of early visual pathways but can also arise from perturbations of higher-order cognitive mechanisms.

Consistent with this idea, our findings further demonstrate that manipulating the contents of working memory, which likely involves higher-order cognitive processes, can induce a similar behavioural dissociation in healthy human observers. Thus, the present paradigm may provide a rigorous way for future neuroscientific studies to investigate the neural basis of conscious perception in nonclinical observers. Unlike classic methodologies, such as masking, which generally reduce processing capacities as well as subjective perception (e.g., Breitmeyer, 2008; Knotts et al., 2018), the present paradigm, by selectively modulating subjective aspects of perception, may help identify neural processes associated with conscious perception while minimizing confounds related to processing capacity.

One may argue that the selective modulation of Y/N sensitivity is not because it is more subjective, but simply because it was naturally harder than 2-IFC tasks (e.g., Macmillan & Creelman, 1991). In other words, working memory may selectively influence tasks that are sufficiently demanding and impose a higher processing load. However, 2-IFC performance was matched using the same staircase procedure, and therefore Y/N performance also remained within a comparable performance range across experiments. Therefore, task difficulty alone cannot explain why the selective impairment disappeared when the Y/N task was formed as a discrimination task (Experiment 2).

A closely related concept to working memory is mental imagery (Tong, 2013; Keogh & Pearson, 2011), which also shares neural substrates in the early visual cortex with visual perception (Dijkstra et al., 2017; Albers et al., 2013). One possibility is that participants maintained the working memory items in an imagery-like format. If so, the observed interaction between working memory and perception may partly reflect mechanisms similar to those reported for visual imagery, which has been shown to boost detection of congruent near-threshold stimuli (Dijkstra et al., 2021). However, working memory content is what we actually perceived in the recent past, whereas imagery could also refer to future and hypothetical situations. This unique relationship with perception and working memory makes it interesting to investigate how they interact and how they are distinguished from each other (Teng & Kravitz, 2019; Lau, 2022). Future studies will therefore be needed to determine whether the interaction observed here reflects imagery-related mechanisms or instead depends on processes that are special for working memory representations.

One limitation of this study may be that we investigated the relationship between working memory and perception using only face and scene stimuli. These distinctive categories may have different perceptual properties, and whether the selective modulation of subjective detection can be generalized to other visual inputs, such as motion, orientation, or colour, remains unknown. Future studies should examine whether similar effects can be generalized across the broader range of visual stimuli.

In conclusion, the current study revealed that what we are holding in mind can change what we subjectively perceive. Holding incongruent content in working memory relatively interrupted the sensitivity of concurrent detection even when discrimination performance was kept constant. Here, we propose a novel psychophysical paradigm for inducing relative blindsight-like behavior in nonclinical observers.

## Declaration of interests

### Competing interests

The authors declare no competing interests.

## Data availability

The behavioural data from all the experiments are publicly available via [https://github.com/wonyi12/Blindsight_WM/tree/main].

## Code availability

The analysis codes are publicly available via [https://github.com/wonyi12/Blindsight_WM/tree/main].

## Author information

### Contributions

W.C, T.N, and H.L designed research. W.C performed experiments and data analyses. W.C, T.N, and H.L wrote the manuscript.

### Corresponding authors

Correspondence to Wonyi Che or Tomoya Nakamura.

## Notes

### Competing Interest Statement

The authors have declared no competing interest.

### Summary of Updates

This version contains the final revisions to the manuscript.

