## Supplementary material for "Relative Blindsight Induced by Incongruency with Working Memory": Fig S1, S2, S3

**Preliminary Experiment**

In a preliminary experiment, a total of 17 university students (11 females, 6 males, mean age = 22.9 years, SD = 2.5) participated in a preliminary experiment. One participant was excluded based on pre-registered criteria (i.e., HR < FAR at least in one condition), so data from 16 participants were used for analysis. The eligibility criteria and ethical procedures were identical to those in Experiment 1.

We initially collected data from eight pilot participants and used these data to estimate the sensitivity ratio (measure mentioned below). A simulation-based power analysis for a one-way repeated-measures ANOVA (Lakens & Caldwell, 2021) indicated that a total sample size of 16 participants would provide >80% power to detect the difference in sensitivity ratio across conditions at α = .05 (for more details, see the Power Analysis section below). For a preliminary experiment, we report data from 8 pilots and 8 new participants. The pilot data were reported together with the final sample size plan as a preprint (https://www.biorxiv.org/content/10.1101/2025.11.14.688399v2.full).

The participants performed the same working memory task accompanied by perceptual tasks during the delay. Unlike in Experiment 1, however, a blank was replaced with a scrambled face interval in the 2-IFC task. In contrast, the Y/N task required the judgment of stimulus presence vs. absence (i.e., face vs blank).

The scaling 2-IFC sensitivity by √2 was comparable to Y/N sensitivity (Fig S1a; SC: scaled 2-IFC $d_a$ = 1.212, Y/N $d_a$ = 1.206; CC: scaled 2-IFC $d_a$ = 1.224, Y/N $d_a$ = 1.318; CI: scaled 2-IFC $d_a$ = 1.202, Y/N $d_a$ = 0.844). After scaling, the task effect was not significant ($F$(1,15) = .326, $p$ = .576, $\eta^2_G$ = .008), indicating that the canonical √2 relationship between Y/N and 2-IFC sensitivity was preserved overall.

The detection sensitivity ($d_a$) ratio differed significantly across congruence conditions ($F$(2,30) = 3.644, $p$ = .038, $\eta^2_G$ = .072). There was a significant difference between the CC and CI ($t$(15) = 3.034, $p$ = .008, Hedges' $g$ = .627), whereas the other pairwise comparisons were not significant (SC–CC: $t$(15) = −0.530, $p$ = .604, Hedges' $g$ = −.143; SC–CI: $t$(15) = 1.917, $p$ = .074, Hedges' $g$ = .448).

To see why Y/N sensitivity selectively drops in the CI condition, we compared hit rate (HR) and false alarm rate (FAR) separately across conditions (Fig S1b). HR showed a significant effect of congruence ($F$(2,30) = 21.141, $p$ < .001, $\eta^2_G$ = .170), whereas FAR did not ($F$(2,30) = 2.608, $p$ = .090, $\eta^2_G$ = *.005*). Pairwise comparisons indicated that HR was significantly reduced in the CI condition relative to both SC and CC (SC–CI: $t$(15) = 5.273, $p$ < .001, Hedges' $g$ = .966; CC–CI: $t$(15) = 5.715, $p$ < .001, Hedges' $g$ = .867). These findings suggest that the reduction in Y/N sensitivity observed in the CI condition was primarily driven by the failure to notice the stimulus.

Since we matched 2-IFC performance by manipulating stimulus contrast, one possibility was that differences in the contrast used accounted for the reduced Y/N sensitivity in the CI condition. However, this was not the case. The contrast thresholds did not differ across the conditions ($F(2,30) = 3.140$, $p = .058$, $\eta_G^2 = .020$). Thus, the reduced Y/N sensitivity in the CI condition cannot be explained by differences in stimulus intensity.

Overall, the preliminary experiment also provided evidence that maintaining category incongruent information in working memory selectively impaired Y/N performance when general perceptual capacity was controlled. Consistent with Experiment 1, working memory could selectively modulate Y/N sensitivity when the task required pure face detection. However, because the stimulus alternatives differed between the two perceptual tasks (face vs scrambled face in 2-IFC; face vs blank in Y/N) in the preliminary experiment, it remained unclear whether this selective impairment reflected the pure detection nature of the Y/N task or differences in the stimulus alternatives themselves. Experiment 1 addressed this ambiguity by matching the stimulus alternatives across tasks while preserving pure detection in Y/N and replicated the selective impairment under the category-incongruent condition. Together with the absence of this effect when Y/N required discrimination (face vs scrambled face) in Experiment 2, we conclude that the selective drop in Y/N sensitivity occurs specifically when the Y/N task constitutes a pure detection paradigm, irrespective of the particular implementation of processing capacity matching.

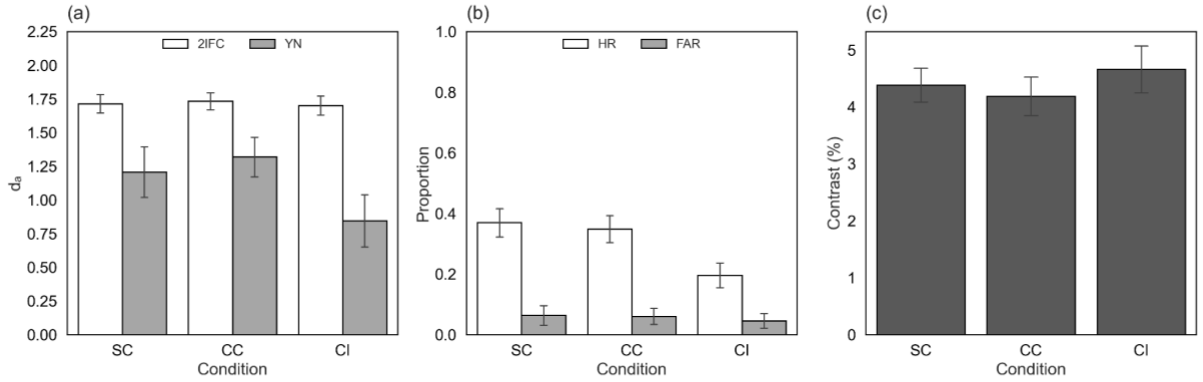


***Fig S1. Results of Preliminary experiment.*** *(a) Sensitivity ($d_a$) in the 2-IFC and Y/N tasks. (b) Hit Rate (HR) and False Alarm Rate (FAR) in the Y/N task. (c) Contrast thresholds derived from staircases in 2-IFC tasks. Bar graphs indicate inter-individual averages, with error bars indicating standard errors of the mean.*

### Additional Analyses

**Power analysis.** We performed a power analysis to determine the sample size using the pilot data (i.e., the initial half 8 of the participants in a preliminary experiment). We focused on the sensitivity ratio (Y/N $d_a$ / 2-IFC $d_a$) as the dependent measure of our primary interest. Assuming the common standard deviation (SD = 0.34, pooled among conditions in pilot) and repeated-measures correlation (R = .74, averaged among pairs of conditions in pilot), we simulated the power of detecting the difference in sensitivity ratio we observed from 8 pilot participants (SC = 0.41, CC = 0.44, CI = 0.25) by one-way repeated-measures ANOVA (Lakens & Caldwell, 2021). We found that at least 16 participants would be necessary to have $1-\beta > .80$ to detect the observed effect at $\alpha = .05$.

**Analysis of metacognitive ability.** We quantified metacognitive sensitivity, i,e, the degree to which participants discriminated between their own correct and incorrect decisions, following the meta-SDT framework (Maniscalco & Lau, 2012). Instead of collecting participants' reported confidence levels, we inferred confidence from reaction times (RT), analogous to our approach in deriving type-1 zROC. We divided trials into three confidence bins (high, medium, and low), each for correct and incorrect responses. Metacognitive sensitivity ($meta\ d_a$) was estimated as $d_a$, which produces this type-2 contingency table with maximum likelihood (Maniscalco & Lau, 2012). The signal-to-noise ratio $s$ was fixed at the value derived from type-1 zROC analysis (mentioned above) when fitting the meta-SDT model, to make the detection and metacognitive sensitivities comparable.

Lastly, metacognitive sensitivity was normalized by detection sensitivity to obtain metacognitive efficiency:

$$M_{ratio} = \frac{meta\ d_a}{d_a}$$

$M_{ratio}$ reflects the extent to which metacognitive performance approaches the ideal level ($M_{ratio} = 1$) predicted from the individual's perceptual sensitivity. We presented it in Fig S2.

**Experiment 1.** In the 2-IFC task (Fig S2a), mean meta-$d_a$ ranged from 1.11 to 1.34 across conditions, whereas mean M-ratio ranged from 0.73 to 0.85 (Fig S2b). To examine whether working memory congruency influenced metacognitive performance, we compared both meta-$d_a$ and m-ratio across the three congruency conditions. Neither meta-$d_a$ ($F(2,30) = 1.740$, $p = .193$, $\eta^2_G = .057$) nor m-ratio ($F(2,30) = 1.103$, $p = .345$, $\eta^2_G = .033$) differed significantly across conditions. These findings indicate that working memory congruency did not influence metacognitive sensitivity or metacognitive efficiency in Experiment 1.

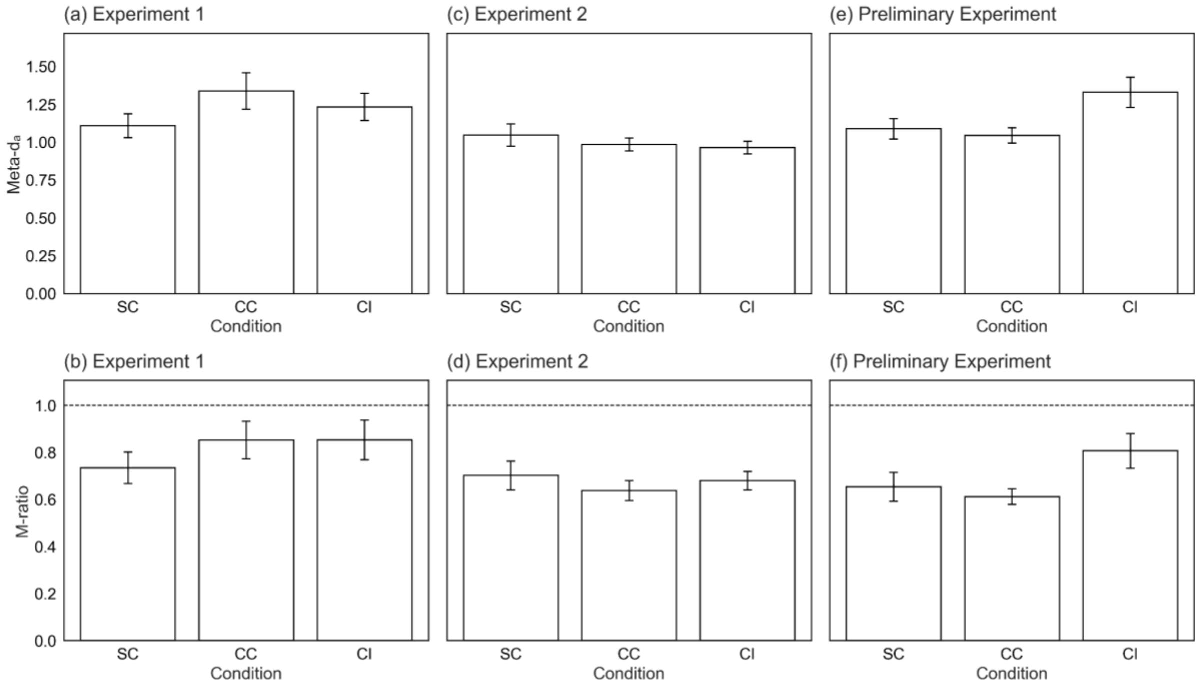


***Fig S2. (a),(c),(e) Metacognitive sensitivity in the 2-IFC task, (b),(d),(f) M-ratio in the 2-IFC task, assuming the unequal variance model.*** *Beyond the detection sensitivity, we explored whether working memory influences metacognitive sensitivity (meta-$d_a$) using reaction time data from the detection task (Miyoshi et al., 2026). Meta-$d_a$ is the participants' ability to monitor the accuracy of their perceptual decisions (Maniscalco & Lau, 2012). Error bars represent SEM across participants. The dashed lines in panels b, d, and f indicate an M-ratio of 1.*

**Experiment 2.** In the 2-IFC task (Fig S2c), mean meta-$d_a$ ranged from 0.96 to 1.05 across the three congruency conditions, whereas mean M-ratio ranged from 0.64 to 0.70 (Fig S2d). Repeated-measures ANOVA revealed that neither meta-$d_a$ ($F(2,30) = 1.247$, $p = .302$, $\eta_G^2 = .027$) nor M-ratio ($F(2,30) = 0.902$, $p = .416$, $\eta_G^2 = .020$) differed significantly across conditions. Therefore, working memory congruency did not alter either metacognitive sensitivity or metacognitive efficiency in Experiment 2.

**Preliminary experiment.** In the 2-IFC task (Fig S2e), mean meta-$d_a$ ranged from 1.05 to 1.33 across conditions, whereas mean M-ratio ranged from 0.61 to 0.81 (Fig S2f). Repeated-measures ANOVA revealed a significant effect of working memory congruence on both meta-$d_a$ ($F(1.306, 19.592) = 4.502$, $p = .038$, $\eta_G^2 = .155$), following Greenhouse–Geisser correction for violation of sphericity. However, the M-ratio was not significant after the correction ($F(1.430, 21.456) = 3.188$, $p = .075$, $\eta_G^2 = .120$). Bonferroni-corrected pairwise comparisons showed that meta-$d_a$ was significantly higher in the CI than in the CC condition ($t(15) = 2.872$, $p = .035$, Hedges' $g = .879$), whereas the remaining comparisons were not significant.

### Additional ANOVAs

**Experiment 1.** For the working memory task, accuracy was well above chance (25%). Still, it did not reach ceiling levels regardless of the congruence condition or the perceptual task (2-IFC or Y/N) performed in the delay (Fig S3a), confirming that the participants understood and followed the task instructions reasonably well while maintaining a sufficiently demanding memory load. A two-way repeated-measures ANOVA revealed no significant main effect of task, indicating that there was no difference in working memory accuracy between 2-IFC and Y/N ($F(1,15) = 1.691$, $p = .213$, $\eta_G^2 = .004$). There was also no significant main effect of congruence ($F(2,30) = 0.555$, $p = .580$, $\eta_G^2 = .003$). More importantly, interaction was not significant ($F(2,30) = 2.439$, $p = .104$, $\eta_G^2 = .004$).

For the perception task, accuracy in the 2-IFC task was successfully matched, targeting near ~79% accuracy (SC: 79.0%, CC: 79.6%, and CI: 78.3%). The Y/N accuracy (SC: 67.2%, $t(15) = 12.003$, $p < .001$; CC: 63.3%, $t(15) = 12.369$, $p < .001$; CI: 56.5%, $t(15) = 6.154$, $p < .001$) was above chance. A 2 x 3 repeated-measures ANOVA revealed a significant main effect of task ($F(1,15) = 248.839$, $p < .001$, $\eta_G^2 = .832$), reflecting that the Y/N was significantly lower than the 2-IFC accuracy. There was also a significant main effect of congruence ($F(2,30) = 32.733$, $p < .001$, $\eta_G^2 = .292$) and the interaction ($F(2,30) = 12.000$, $p < .001$, $\eta_G^2 = .234$; Fig S3b).

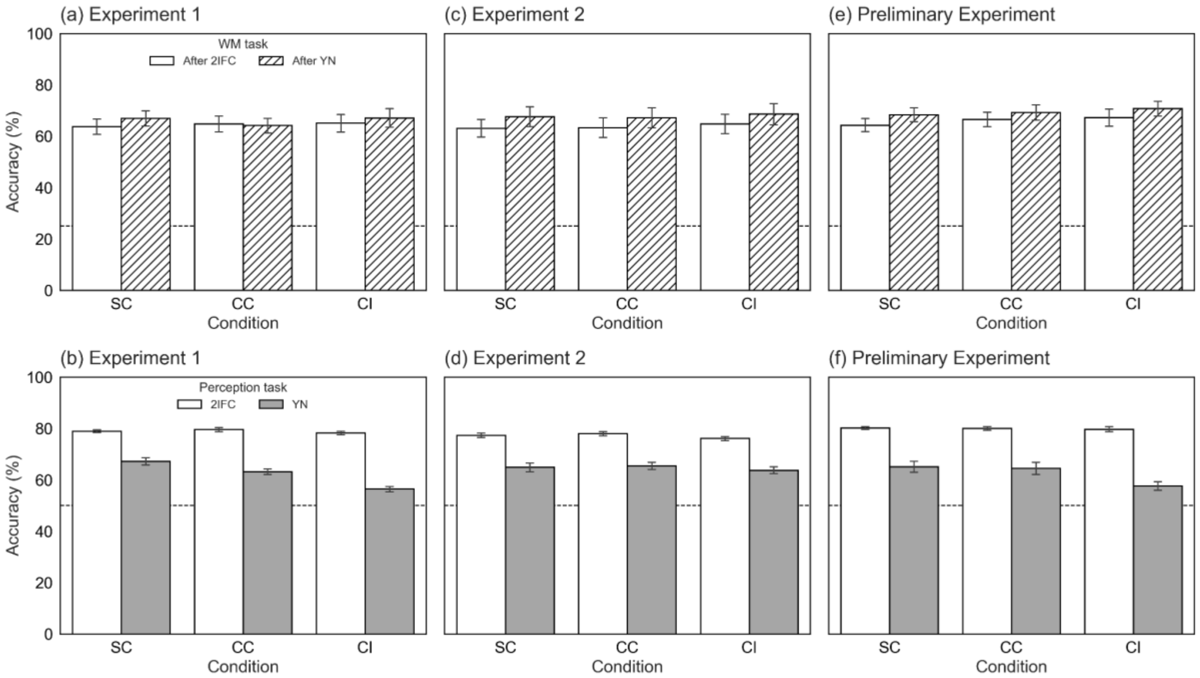


***Fig S3. (a),(c),(e) Accuracies (%) for the working memory task for Experiments 1,2, and preliminary Experiment, (b),(d),(f) Accuracies for the 2-IFC and Y/N task.*** *Gray dashed lines indicate chance performance (25% for the working memory task; 50% for the 2-IFC and Y/N tasks). Error bars indicate standard errors of the mean.*

Canonical √2 relationship between 2-IFC and Y/N was preserved across congruence, as there was no significant effect of task ($F$(1,15) = 0.248, $p$ = .626, $\eta_G^2$ = .004). However, the main effect of congruence ($F$(2,30) = 8.419, $p$ = .001, $\eta_G^2$ = .122) and interaction ($F$(2,30) = 5.728, $p$ = .008, $\eta_G^2$ = .090) were significant. Separate analyses within each congruence showed no significant difference between Y/N and scaled 2-IFC sensitivity in either the SC condition ($t$(15) = 2.198, $p$ = .132, Hedges' $g$ = 0.771) or the CC condition ($t$(15) = -0.356, $p$ = 1.000, Hedges' $g$ = -0.123). In contrast, the CI condition showed significantly lower Y/N than scaled 2-IFC sensitivity, suggesting the largest deviation from the √2 calculation ($t$(15) = -2.954, $p$ = .030, Hedges' $g$ = -0.961). These findings suggest that the canonical √2 relationship between discrimination and detection sensitivity may be selectively weakened under category-incongruent working memory conditions.

**Experiment 2.** For the working memory task, accuracy was well above chance (25%), and it still did not reach ceiling levels (Fig S3c). This confirms that the participants understood and followed the task instructions reasonably well. The memory load was high enough to see the potential influence on the perceptual task. A two-way repeated-measures ANOVA revealed that working memory accuracy was higher after Y/N trials than after 2-IFC trials ($F$(1,15) = 8.323, $p$ = .011, $\eta_G^2$ = .019). Although working memory accuracy was numerically highest in the category-incongruent condition, there was no significant main effect of congruence ($F$(2,30) = 0.610, $p$ = .550, $\eta_G^2$ = .002), nor the interaction, $F$(2,30) = 0.089, $p$ = .915, $\eta_G^2$ < .001.

For the perception task, the accuracies of the 2-IFC and Y/N tasks were separately shown in Fig S3d. We calibrated the stimulus contrast in the 2-IFC task targeting ~79% accuracy, and mean accuracy was actually matched across congruences as we expected (SC: 77.4%, CC: 78%, and CI: 76.1%). The Y/N accuracy (SC: 64.9%, CC: 65.5%, and CI: 63.8%) was above chance but lower than the 2-IFC accuracy ($F$(1,15) = 148.685, $p$ < .001, $\eta_G^2$ = .637). However, neither the main effect of congruence ($F$(2,30) = 1.442, $p$ = .252, $\eta_G^2$ = .025) nor the task × congruence interaction ($F$(2,30) = 0.007, $p$ = .993, $\eta_G^2$ < .001) was significant.

Consistently, the 2-IFC sensitivity scaled by √2 was also comparable to Y/N sensitivity. There was no main effect of task ($F$(1,15) = 0.183, $p$ = .675, $\eta_G^2$ = .002), congruence ($F$(2,30) = 1.198, $p$ = .316, $\eta_G^2$ = .026), or the interaction ($F$(2,30) = 0.278, $p$ = .759, $\eta_G^2$ = .004).

**Preliminary experiment.** For the working memory task, overall accuracy was well above chance (> 25%) but did not reach ceiling levels (Fig S3e). This confirms that the participants understood and followed the task instructions reasonably well, while the memory load was high enough to see the potential influence on the perceptual task. A two-way repeated-measures ANOVA revealed that working memory accuracy was higher when accompanied by Y/N than when accompanied by 2-IFC ($F$(1,15) = 8.825, $p$ = .010, $\eta_G^2$ = .023). The main effect of congruency was not significant ($F$(2,30) = 3.069, $p$ = .061, $\eta_G^2$ = .009). More importantly, the task-by-congruence interaction was not significant ($F$(2,30) = .257, $p$ = .775, $\eta_G^2$ = .001).

For the perception task, accuracy in the 2-IFC task was successfully matched around 79% across congruence conditions through staircasing, as intended (SC: 80.2%, CC: 80.1%, and CI: 79.8%). The accuracy in the Y/N task was above chance (SC: 65.2%, $t$(15) = 7.154, $p$ < .001; CC: 64.5%, $t$(15) = 6.187, $p$ < .001; CI: 57.7%, $t$(15) = 4.533, $p$ < .001). A 2 x 3 repeated-measures ANOVA revealed a significant main effect of the task ($F$(1,15) = 65.353, $p$ < .001, $\eta_G^2$ = .679), reflecting lower accuracy in the Y/N task than the 2-IFC task. The main effect of congruence ($F$(2,30) = 13.681, $p$ < .001, $\eta_G^2$ = .081) and the interaction ($F$(2,30) = 8.353, $p$ = .001, $\eta_G^2$ = .066) were also significant (Fig S3f).

To further examine whether the canonical √2 relationship between 2-IFC and Y/N was preserved across congruence, we compared Y/N sensitivity with √2 -scaled 2-IFC sensitivity. The main effect of the task was not significant ($F$(1,15) = .326, $p$ = .576, $\eta_G^2$ = .008), consistent with the theoretical √2 relationship between the two tasks. However, the main effect of congruence, remained, $F$(2,30) = 4.952, $p$ = .014, $\eta_G^2$ = .042, and the interaction was also significant, $F$(2,30) = 3.387, $p$ = .047, $\eta_G^2$ = .036. Follow-up paired comparisons revealed no significant difference between Y/N and √2-scaled 2-IFC sensitivity in any congruence condition (SC: $t$(15) = −0.030, $p$ = 1.000, Hedges' $g$ = −0.011; CC: $t$(15) = 0.579, $p$ = 1.000, Hedges' $g$ = 0.209; CI: $t$(15) = −1.747, $p$ = .303, Hedges' $g$ = −0.614).
